# *Vespula pensylvanica* locate odor sources across diverse natural wind conditions

**DOI:** 10.1101/2025.10.01.679804

**Authors:** Jaleesa Houle, Floris van Breugel

**Affiliations:** Dept. of Mechanical Engineering, University of Nevada, Reno; Integrative Neuroscience Program, University of Nevada, Reno

**Keywords:** olfactory navigation, natural wind, olfactory search

## Abstract

Many organisms across ecosystems track odor plumes to locate mates and food. In flying insects, the task of localizing an odor source is particularly challenging due to the complicated dynamics associated with wind flow and odor plume dispersion through spatially complex environments. Although wind tunnel experiments have been instrumental for answering many questions related to olfactory search, such experiments cannot replicate the complexity of natural wind conditions. Thus, our knowledge of how real-world wind characteristics influence insects’ success and strategies to locate odor sources remains an open area of investigation. Here, we tested whether certain wind conditions were more favorable for foraging insects by comparing yellowjacket arrival times and corresponding wind conditions across three distinct natural environments. Our results indicate that *Vespula pensylvanica* are capable of locating odor sources across the full range of observed wind conditions, without any clear preferences. This suggests that insects have adapted strategies to perform odor localization tasks across the full spectrum of natural wind that they may encounter. Our field-based approach provides insight into key considerations for future wind tunnel experiments which seek to better resolve insect plume tracking in understudied flow regimes.

## INTRODUCTION

Across ecosystems and taxa, many animals rely on olfactory cues to locate food and mates [27, 35, 36, 23]. For flying insects such as pollinators, this task is critical for both their ecology and human agriculture [18]. Despite more than a century of research on insect foraging [4], most of the insights into olfactory search in flight stem from controlled wind tunnel studies. Although invaluable for isolating specific mechanisms, such experiments cannot replicate the complex multiscale turbulence and rapid spatiotemporal variability that characterize natural wind flow [11, 25, 14, 31]. Consequently, our knowledge of how insects perform olfactory search tasks in natural environments remains limited. Although some studies have found that certain wind conditions can be energetically costly and may reduce foraging behaviors [6, 13], others have found that turbulent plume structures may enhance an insect’s ability to locate odor sources [7, 33]. These findings raise the question: is there a favorable range of wind conditions in which insects are more likely to engage in, and succeed at, plume tracking tasks?

Near-surface wind conditions can largely be captured by two correlated metrics [14]: turbulence intensity *T*_*i*_ (the standard deviation of wind speed normalized by the mean), and directional variability *σ*_*θ*_ (the standard deviation of wind direction). Tracking an odor plume across the range of these metrics involves several trade-offs. High mean wind speeds are often associated with lower turbulence intensity (though the reverse is not always true) [14, 7]. In laminar flow, both *T*_*i*_ and *σ*_*θ*_ will be low, resulting in odor plumes with a ribbon-like shape [24]. Conversely, turbulent flow can be characterized by high *T*_*i*_ and *σ*_*θ*_, resulting in highly fragmented and dispersed plumes. Although the ribbon-like plumes generated in higher mean wind speeds would facilitate well established plume tracking algorithms such as the surge-and-cast algorithm [34, 3], such conditions can be difficult for foragers to maneuver. Thus, it is not surprising that at high wind speeds (≥ 3.5 m/s) honey bees reduce flower visitation [13], and orchid bees increase energy expenditures [6]. Meanwhile, some level of directional variability (and subsequently lower wind speeds) has been hypothesized to help with plume tracking due to enhanced plume dispersion, which may improve the probability of encountering odor packets [8, 5, 30]. Higher directional variability has been shown to improve odor source localization for flying moths [33], and despite the energetic disadvantages caused by wind disturbances [29], outdoor observations of orchid bees show that turbulence can increase visitation rates [8]. Even over large spatial scales spanning up to 1 km, release and recapture experiments have shown that fruit flies are capable of locating bait traps across a wide range of wind conditions [20]. Taken together, these studies provide evidence that insects may have adapted strategies that allow them to locate odor sources over a variety of wind conditions, but suggest that certain conditions, such as high wind speeds, may be less favorable.

One of the challenges with natural wind is that its behavior can change over relatively short periods of time. Thus, testing the hypothesis that certain natural wind conditions are more favorable for successful source localization requires quantifying the arrival times of a single species across a range of wind conditions with a fine temporal resolution. To perform such an experiment we chose to study a relatively ubiquitous neighbor of pollinators, yellowjackets (*Vespula spp*.). Although most yellowjacket species are social, evidence suggests that unlike bees neither distance nor directional information is shared among nestmates [15, 16], making them a strong candidate for plume tracking studies across different environments. We tested whether certain wind conditions were favorable for plume tracking by deploying baited yellowjacket traps across three distinct outdoor environments and tracking yellowjacket arrivals alongside detailed wind measurements. We compared wind characteristics preceding arrival events to randomly sampled measurements across the period of study. Our findings indicate that yellowjackets are able to locate odors across turbulent and high-speed wind conditions, suggesting robust plume tracking capabilities.

## MATERIALS AND METHODS

### Site locations and collection protocol

Yellowjacket arrivals at baited traps (Figure 1A) were tracked between June-October 2023 in three distinct environments (Figure 1B). The commercially available and reusable yellowjacket traps (*RESCUE!*©) were modified with photo interrupters (SparkFun, GP1A57HRJ00F) at each entrance hole (Figure 1A). A real-time clock (RTC DS3231) was installed on an Adafruit Feather M0 Adalogger microcontroller to record arrival times of yellowjackets entering the trap (Figure 1A). A total of six modified yellowjacket traps were deployed, with two in each environment, placed roughly 10 meters apart. Since photo interrupters are light sensitive, we found during our initial launch that some of the traps placed in full-sun areas would not recognize trigger events. To fix this, we wrapped the traps in aluminum foil, which shielded the photo interrupters from sun exposure. In addition to these six modified traps, a separate unmodified trap was placed nearby as a control to verify that our modifications did not influence the ability of insects to successfully enter the traps.

**Figure 1.**
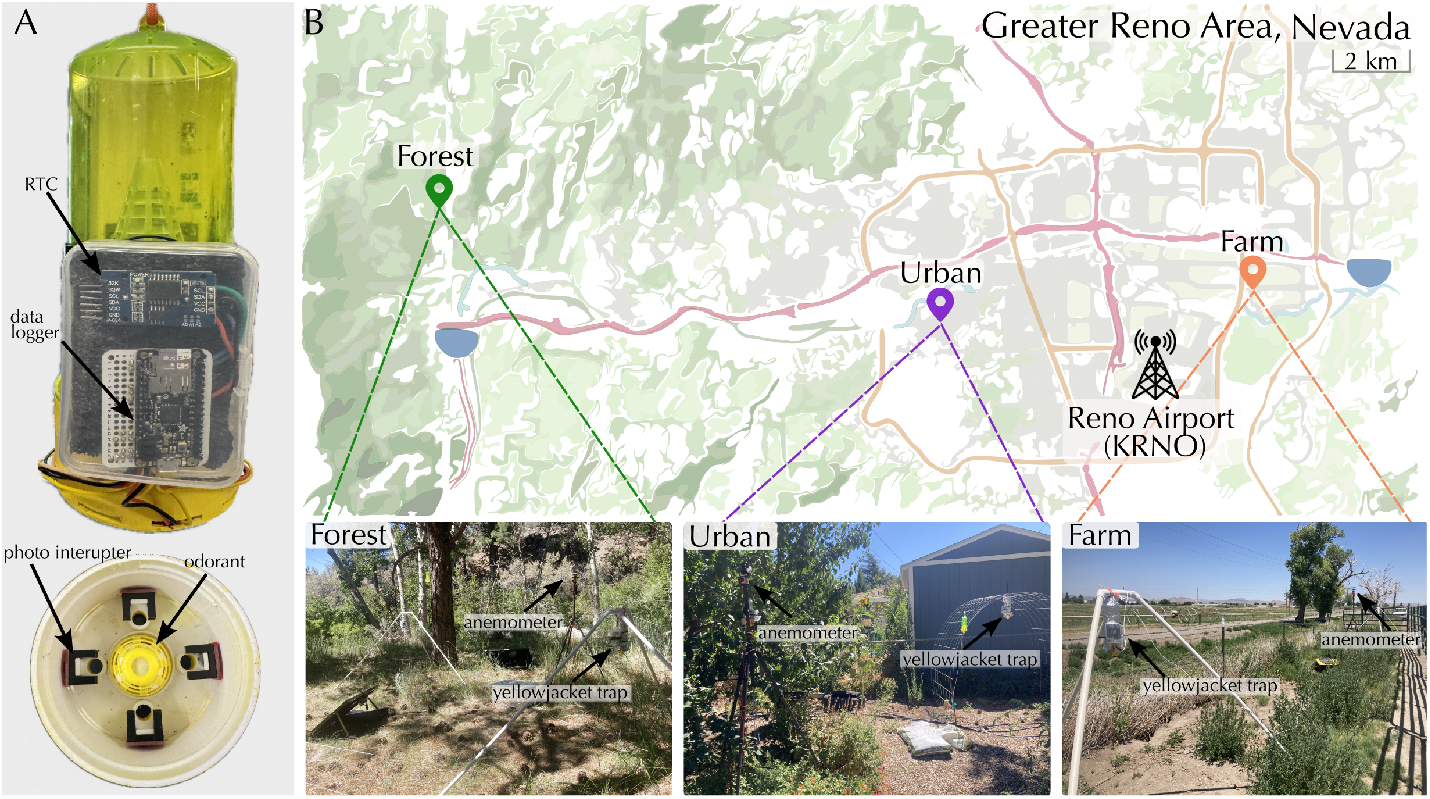
Experimental setup and site locations. (A) Modified yellowjacket traps equipped with a datalogger, real-time clock (RTC), and photo interrupters at each entrance hole. (B) Approximate site locations and pictures of each site.

At the farm and forest sites, solar panels and external batteries were used to keep all equipment powered. The urban site was powered through long extension cords to reach standard electrical outlets. There were instances in which power was lost due to various weather events and wildlife interference. To minimize these occurrences, sites were visited twice a week. Even still, this amount of site visitation was not enough to fully mitigate data loss. Overall, across the 4 months of deployment, wind data was successfully recorded across a total of approximately 2 months worth of unique days.

Prior research has found that yellowjackets may forage as far as 3,100 feet away from their nest [1]. Due to limited land use permissions, we were not able to canvas these distances in the surrounding areas. Although such information would be beneficial for better understanding the relative populations and success rates of foragers across environments, we have limited the scope of our analysis solely to intra-class comparisons and make no speculation on overall foraging success rates.

### Odorant Selection and Maintenance Protocol

Before starting this experiment, several traps were set up in our candidate environments to verify that populations existed in all locations. Yellowjackets were successfully captured in all environments, although the relative abundance varied. During these preliminary tests, both heptyl butyrate and meat (pork) were tested as potential lure attractants. Heptyl butyrate is the standard chemical used in *RESCUE!* yellowjacket traps, and has been a known yellowjacket attractant since the late 1960s [9, 21]. This compound has been found to be particularly effective in the western US, where *Vespula pensylvanica* is a common pest [2, 24]. Our preliminary tests found that traps with meat contained a considerable amount of fly bycatch, whereas the traps with heptyl butyrate caught only *Vespula pensylvanica*. As such, we chose to proceed with heptyl butyrate in our experiment.

In an effort to emit a relatively consistent odor concentration, *RESCUE!* Yellowjacket Attractant Cartridges (model YJTC) were utilized for each trap (Figure 1A). These cartridges weigh approximately 0.4 ounces and are made up by weight of 63.9% heptyl butyrate and 36.1% polyethylene terephthalate (inert polyester fibers). The combination of liquid heptyl butyrate soaked in polyester fibers allows these cartridges to remain effective for up to 10 weeks (based on manufacturer claims). To further maintain a consistent release of odorant concentration throughout the duration of our experiment, cartridges were routinely replaced every 4-5 weeks.

### Species identification

The traps used for our experiments were designed such that yellowjackets could not escape after entering the trap. This allowed us to verify that the amount of trigger events and corresponding insect remains were consistent throughout the collection period. The collected specimens were identified by submitting pictures of several samples to online community identification tools, *iNaturalist* and *BugGuide*. Several specialists identified our samples as *Vespula pensylvanica*, i.e. the Western yellowjacket.

### Wind collection

In addition to recording insect arrival times, wind recordings were taken using a 3D ultrasonic anemometer (Trisonica™Mini, LI-COR Environmental) at a rate of 10Hz at each site. The Trisonica™Mini has a specified accuracy of ±0.2 m/s in wind speeds between 0-10 m/s, ±.2% m/s in wind speeds between 11-30 m/s, ±2^*°*^C for temperature, and ±1° for horizontal wind direction. Additionally, the Mini is equipped with a pressure sensor that operates within a range of 50-115 kPa, resolution of 0.1 kPa and accuracy of *±*1 kPa. Note that our sensors were not configured to record air pressure, but instead air density, which is derived from the speed of sound and air pressure readings. Using the ideal gas law, we were able to back-calculate air pressure such that *P* = *ρRT* where *ρ* is the estimated air density (*kg/m*^3^), *R* = 287.06*J/*(*kgK*), and *T* is the air temperature. Each anemometer was manually leveled and oriented toward North, and placed roughly 2 meters above ground level. Teensy 3.5 controllers with microSD cards were used for data-logging while a separate GPS receiver (GP-20U7 (56 Channel), Sparkfun) recorded time and location information.

### Data analysis and statistics

To compare variability in wind speed across days and environments, we used the non-dimensional metric for turbulence intensity, defined as *T*_*i*_ = *σ*_*h*_*/U*_*h*_, where *σ*_*h*_ is the standard deviation of horizontal wind speed and *U*_*h*_ is the mean horizontal wind direction. We computed the standard deviation in wind direction, *σ*_*θ*_, to assess directional variability. We calculated *T*_*i*_ and *σ*_*θ*_ for the 5 minutes leading up to each trigger, and for randomly sampled 5 minute wind measurements across all days in which wind data were successfully recorded (i.e. “null” wind events). In total, there were 55 unique days of wind data for the urban site, 57 for the farm, and 59 for the forest. Null events were selected by randomly choosing start indexes across the datasets, up to a maximum of 30 samples per day. Time segments were selected such that there was no overlap between individual samples. We then performed 2-D Kolmogorov–Smirnov (KS) tests individually for each environment to assess whether the *T*_*i*_ and *σ*_*θ*_ observed prior to insect arrivals had the same distribution as randomly sampled null wind events.

Since yellowjackets primarily arrived during daytime hours, and wind regimes are known to transition from night and daytime [22], we limited our random null sampling to measurements between 6am to 8pm local time. Additionally, since temperature is a known confounding factor that can influence both wind conditions and insect foraging activities [11, 12], we limited the sampled null wind events to be within the minimum and maximum temperatures recorded during arrival events of each respective environment. In addition to these KS tests, we performed a non-parametric Mann-Whitney U test to assess the likelihood that the wind speeds observed preceding arrival events at the farm site were drawn from the same distribution as the randomly sampled null wind events. This test was only relevant for the farm site due to the much larger range of wind speeds observed.

### Controlling for additional atmospheric variables

It is generally expected that quick changes in barometric pressure can precede adverse weather events, which subsequently impact insect activity [26, 10, 19]. To verify that atmospheric pressure fluctuations did not influence our results, we compared the standard deviation in pressure across all 5 minute null wind and trigger events (Supplemental Figure S1A), where large standard deviations would indicate larger fluctuations.

Prior studies have also shown that rainfall can impact foraging behavior of yellowjackets [17]. To account for this, we retrieved daily precipitation records from the National Oceanic and Atmospheric Administration’s (NOAA) National Center for Environmental Information (NCEI) at the Reno Tahoe Airport weather station (KRNO, Figure 1B) for all days of data collection. In total, there were 10 days in which precipitation was reported, and trigger events occurred on 6 of those days. Only two of those dates reported more than 0.1 inches of total rain, with half of them reporting between 0.01-0.03 inches (Supplemental Table S1). Since these daily totals do not give insight into the time of day, duration, or intensity of storms, we compared *T*_*i*_ and *σ*_*θ*_ while excluding all days in which any precipitation occurred, in addition to excluding null wind samples with standard deviations in pressure that were higher than the maximum observed during trigger events (Supplemental Figure S1B-C).

## RESULTS

Across the four months of data collection, we recorded a total of 52 yellowjacket arrivals at the farm site, 224 at the forest site, and 114 at the urban site. Most arrivals occurred during daytime hours between 6am and 8pm (Figure 2A), though there were a few outliers. Unfortunately, power loss due to weather events and anemometer recording errors reduced the quantity of trigger events that had corresponding wind measurements. Trigger events with corresponding wind measurements amounted to a total of 20 at the farm site, 75 at the forest site, and 45 at the urban site. Figure 2B shows a 20 minute sample of wind speed measurements, during which two insects arrived at traps in the urban environment. Across environments, the average temperatures for the 5 minutes preceding arrival events were generally between 10^*°*^C and 27^*°*^C (Figure 2C). Mean wind speeds averaged over 5 minute intervals generally did not exceed 2 m/s in the urban and forest environments, whereas wind speeds at the farm site for both arrival events and randomly sampled null wind events reached upwards of 6 m/s at times (Figure 2C).

**Figure 2.**
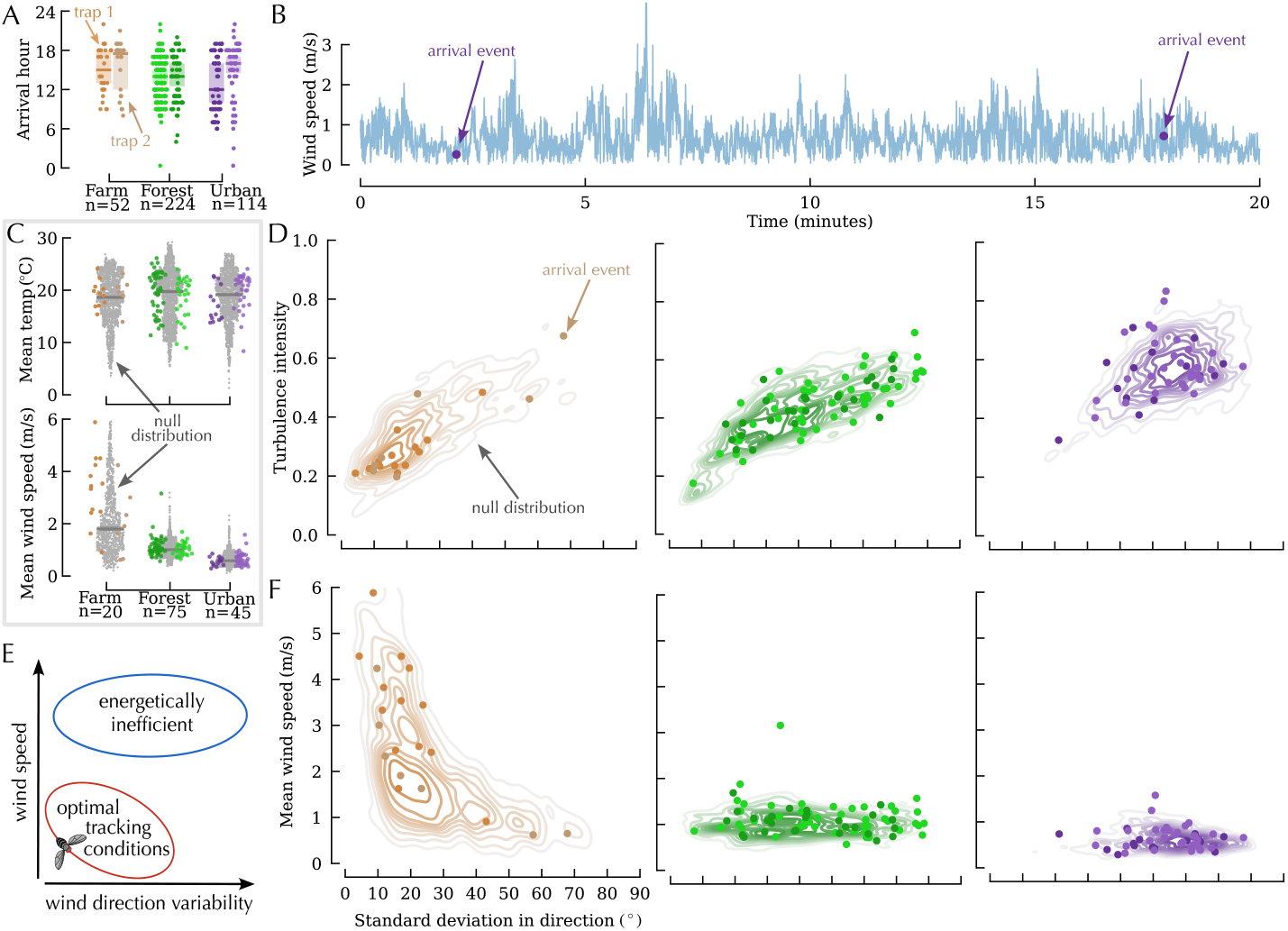
Yellowjackets are capable of foraging across a diverse set of wind conditions. A) Arrival times recorded for all yellowjackets across the four month period of recording. (B) Sample time series for the urban site during which two yellowjackets entered traps. (C) 5 Minute average temperatures and wind speeds. Colored points correspond to arrival events shown in *A* and gray points correspond to randomly sampled 5 minute null events across all daytime wind recordings for the farm (n=1571), forest (n=1754), and urban (n=1589) sites. Note that the number of yellowjacket arrival events is lower than in A due to various recording malfunctions over the two month collection period. (D) Turbulence intensity (*T*_*i*_) versus standard deviation in wind direction (*σ*_*θ*_) for each environment. Scatter points correspond to 5 minute averages preceding yellowjacket arrivals, and contour lines were generated from all randomly sampled 5 minute null wind events between 6am and 8pm and subject to the same temperature ranges observed across arrival events events (as seen in B-C) for the farm (n=1056), forest (n=1593), and urban (n=1380) sites. (E) Hypothesized range of wind conditions where we might expect to see a concentration of arrivals. (F) Mean wind speed versus *σ*_*θ*_ for the farm, forest, and urban environments.

### Yellowjackets arrived across all observed wind conditions

The spread of turbulence intensities (*T*_*i*_) and standard deviations in wind direction (*σ*_*θ*_) varied across environments, with generally lower values of *T*_*i*_ and *σ*_*θ*_ reported at the farm site, and much larger values of *T*_*i*_ and *σ*_*θ*_ and the urban site (Figure 2D). These results reflect the effects of larger surface complexities on wind profiles. A 2-D Kolmogorov–Smirnov (KS) test was performed to assess the likelihood that the 5-minute *T*_*i*_ and *σ*_*θ*_ values corresponding to arrival events were drawn from the same distribution as the null wind samples. The two distributions were not statistically different for the forest (*D*=0.178, *p=0*.*492*) or urban environments (*D*=0.186, *p=0*.*185*). The KS test for the farm site was statistically significant (*D*=0.355, *p=0*.*022*), with a larger proportion of arrivals occurring in wind conditions with lower *T*_*i*_ and *σ*_*θ*_ as compared to the null distribution. However, we note that adjusting for precipitation and fluctuations in atmospheric pressure resulted in this test being statistically insignificant for the farm site (*D*=0.325, *p=0*.*063*, Supplemental Figure S1B), demonstrating that this trend is not strong and may be influenced by both low sample size and additional atmospheric variables.

Based on prior studies looking at the energetic costs of flight in turbulence and at higher wind speeds [13, 33], we predicted that a larger proportion of yellowjackets would arrive at traps in scenarios where wind speeds and standard deviations in wind direction were lower (Figure 2E). However, comparing randomly sampled 5 minute mean wind speeds to the 5 minute mean wind speeds preceding arrival events as a function of *σ*_*θ*_ demonstrated that a majority of arrivals at the farm site occurred in higher wind speeds with lower directional variability, while plenty of arrivals in both the forest and urban settings occurred across a range of *σ*_*θ*_, but at much lower mean wind speeds (Figure 2F). The non-parametric Mann-Whitney U test was performed to determine if the underlying distribution of wind speeds at the farm site during arrival events was the same as observed across random null samples, and indicated that the two populations were significantly different (*U=7517, p=0*.*027*). This statistical significance did not change when taking barometric pressure fluctuations and precipitation into consideration (Supplemental Figure S1C). Taken together, these results indicate that yellowjackets were able to successfully localize odor sources across a broad range of wind conditions, with no obvious aversion to higher wind speeds, larger turbulence intensities, or greater variability in direction, despite potential energetic constraints.

## DISCUSSION

Our results indicate that *Vespula pensylvanica* are able to successfully track odors over a diverse set of wind conditions. Comparing the null wind distributions to wind measurements directly preceding trigger events allowed us to assess whether certain wind conditions were indeed more favorable for active foragers after accounting for separate factors such as mean temperature and time of day. Statistically testing the distribution of *T*_*i*_ and *σ*_*θ*_ between arrivals and null events confirmed that there was no significant change in trapping for certain wind conditions among foragers in the forest and urban sites. We note that there was a statistically significant difference between the null and arrival population mean wind speeds at the farm site, indicating that higher wind speeds were in fact more favorable in that environment, though the small sample size makes it challenging to draw strong conclusions. It is important to note that additional variables, such as barometric pressure and rainfall, can influence insect behavior [19, 28, 26]. For our data, removing all days in which precipitation occurred, along with null wind samples that had high standard deviations in atmospheric pressure, did not strongly impact our conclusions (Supplemental Figure S1).

Noting the overall large difference in mean wind speeds recorded between the farm and forest/urban sites, one may wonder if the smaller population of arrivals at the farm site may, in part, be caused by higher overall average speeds. Since we do not know the starting population size in each environment, we cannot speculate as to how relatively successful yellowjackets were at reaching the traps as a function of the wind they experienced. However, we note that there were still many null samples in which wind speeds were lower, and many of the farm arrivals occurred while mean wind speeds were in fact higher (*U*_*h*_ *>* 2m/s) than seen at the forest and urban sites. These results indicate that higher wind speeds likely did not strongly impact the ability for yellowjackets to locate the traps. An alternative hypothesis for the proportionally smaller amount of arrivals observed at the farm site is that for sites with less surface roughness elements, such as a farm or desert, the lack of features for dispersing odor plumes to interact with at low wind speeds could potentially reduce the overall volume of odor encounters for plume tracking insects, making plume tracking in a simple environment more challenging at lower wind speeds and thus making higher speeds more favorable.

Overall, our findings demonstrate that yellowjackets will successfully participate in foraging activities despite potential mechanical or algorithmic disadvantages that the local wind presents. How might the navigation of a forager differ in strong unidirectional winds as opposed to turbulent or directionally variable winds? Based on the range of wind conditions observed preceding arrival events, we hypothesize that flying insects, such as yellowjackets, must employ different navigational strategies across wind conditions to achieve their tracking goals. Although an increasingly large body of studies have demonstrated clear surge-cast behaviors for flying insects in generally consistent wind directions [34, 3], alternative behaviors in other wind regimes have recently been discovered [32]. As such, future works directed toward recreating a larger variety of wind conditions as seen in natural settings will be helpful in unveiling behaviors beyond the well established surge-cast, and more recently established sink-circle, behaviors observed in wind tunnel studies.

## AUTHOR CONTRIBUTIONS

F.v.B. and J.H. conceived the experimental design. J.H. constructed the traps and wrote the corresponding software. J.H. performed all data collection and analysis. F.v.B. and J.H. wrote the manuscript.

## DATA AVAILABILITY

Data will be made available upon publication.

## ACKNOWLEDGMENTS

We would like to thank Anne Leonard for all of her suggestions and feedback, and Molly Allen for helping with design and construction of our modified traps. We would also like to thank Scott Huber for his help in coordinating access to UNR’s Main Station Field Lab.

