## Supplemental Material for "*Vespula pensylvanica* locate odor sources across diverse natural wind conditions"

### SUPPLEMENTARY MATERIAL

#### Daily Precipitation Records

| Date | Total Precipitation (inches) | # Trigger Events |
|---|---|---|
| Aug 14, 2023 | 0.07 | N/A |
| Aug 18, 2023 | 0.02 | 1 (Forest) |
| Aug 22, 2023 | 0.02 | 2 (Farm) |
| Sept 01, 2023 | 0.11 | N/A |
| Sept 02, 2023 | 0.09 | 1 (Forest) |
| Sept 03, 2023 | 0.34 | 5 (Forest) |
| Sept 21, 2023 | 0.02 | 3 (Forest) |
| Sept 30, 2023 | 0.05 | N/A |
| Oct 01, 2023 | 0.01 | N/A |
| Oct 11, 2023 | 0.03 | 2 (Forest) |

Supplementary Table S1: Dates, total precipitation, number of trigger events, and environment where trigger events occurred across all days in which rain was recorded at the nearby Reno Tahoe Airport (KRNO) weather station shown in Figure 1B. These data were retrieved from NOAA's NCEI database.

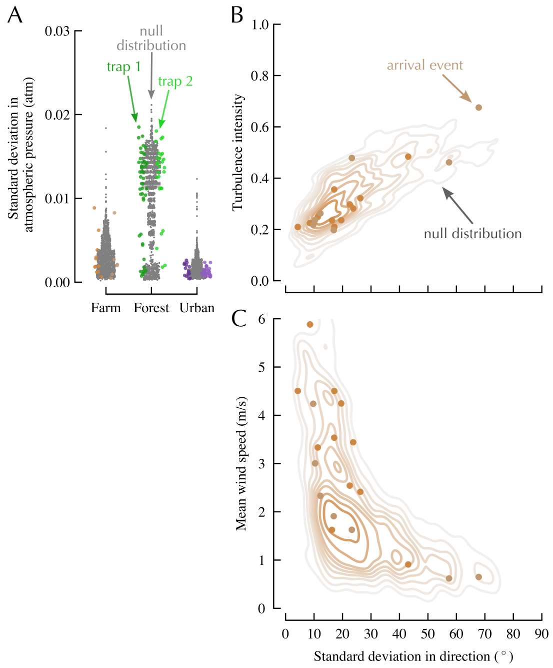


Supplementary Figure S1: **Excluding days with reported precipitation and removing null wind events with large standard deviations in barometric pressure did not strongly impact the relationship between null and trigger event distributions**. (A) Average barometric pressure for randomly sampled 5 minute null events when compared to the 5 minute averages prior to insect arrival events. (B) Turbulence intensity ($T_i$) versus standard deviation in wind direction ($\sigma_\theta$), averaged over 5 minute periods for each environment, excluding the 10 days in which any precipitation was reported at the KRNO weather station. Scatter points correspond to 5 minute averages leading up to yellowjacket arrivals, and contour lines were generated from all randomly sampled 5 minute null wind events between 6am and 8pm and subject to the same temperature and atmospheric pressure ranges observed across arrival events events for the farm (n=1332), forest (n=1530), and urban (n=1489) sites. Computing a 2-D Kolmogorov-Smirnov (KS) test with these adjusted null distributions show that they are not significantly different than the arrival occurances for the farm ($D$=0.325, $P$=0.063), forest ($D$=0.193, $P$=0.153), or urban ($D$=0.16, $P$=0.191) sites. (C) Mean wind speed versus $\sigma_\theta$ for the farm, forest, and urban environments, with scatter points representing 5 minute means preceding arrival events and contour lines corresponding to randomly sampled 5 minute null wind events subject to the same filtering protocol as in *B*. Conducting a Mann Whitney U test on the reduced datasets for mean wind speeds at the farm indicated that the difference in arrival versus null distributions was still statistically significant after adjusting for atmospheric pressure and precipitation ($U$=6385, $p$=0.043).
